# *Heliconius* butterflies host characteristic and phylogenetically structured adult-stage microbiomes

**DOI:** 10.1101/820829

**Authors:** Tobin J. Hammer, Jacob C. Dickerson, W. Owen McMillan, Noah Fierer

## Abstract

Lepidoptera (butterflies and moths) are diverse and ecologically important, yet we know little about how they interact with microbes as adults. Due to metamorphosis, the form and function of their adult-stage microbiomes might be very different from microbiomes in the larval stage (caterpillars). We studied adult-stage microbiomes of *Heliconius* and closely related passion-vine butterflies (Heliconiini), which are an important model system in evolutionary biology. To characterize the structure and dynamics of heliconiine microbiomes, we used field collections of wild butterflies, 16S rRNA gene sequencing, quantitative PCR, and shotgun metagenomics. We found that *Heliconius* harbor simple and abundant bacterial communities that are moderately consistent among conspecific individuals and over time. Heliconiine microbiomes also exhibited a strong signal of host phylogeny, with a major distinction between *Heliconius* and other butterflies. These patterns were largely driven by differing relative abundances of bacterial phylotypes shared among host species and genera, as opposed to the presence or absence of host-specific phylotypes. We suggest that phylogenetic structure in heliconiine microbiomes arises from conserved host traits that differentially filter microbes from the environment. While the relative importance of different traits remains unclear, our data indicate that pollen-feeding (unique to *Heliconius*) is not a primary driver. Using shotgun metagenomics, we also discovered trypanosomatids and microsporidia to be prevalent in butterfly guts, raising the possibility of antagonistic interactions between eukaryotic parasites and co-localized gut bacteria. Our discovery of characteristic and phylogenetically structured microbiomes provides a foundation for tests of adult-stage microbiome function, a poorly understood aspect of lepidopteran biology.

**Importance:** Many insects host microbiomes with important ecological functions. However, the prevalence of this phenomenon is unclear, because in many insect taxa microbiomes have only been studied in part of the life cycle, if at all. A prominent example is the butterflies and moths, in which the composition and functional role of adult-stage microbiomes are largely unknown. We comprehensively characterized microbiomes in adult passion-vine butterflies. Butterfly-associated bacterial communities are generally abundant in guts, consistent within populations, and composed of taxa widely shared among hosts. More closely related butterflies harbor more similar microbiomes, with the most dramatic shift in microbiome composition occurring in tandem with a suite of ecological and life history traits unique to the genus *Heliconius*. Butterflies are also frequently infected with previously undescribed eukaryotic parasites, which may interact with bacteria in important ways. These findings advance our understanding of butterfly biology and of insect-microbe interactions generally.

## Introduction

Insect microbiome research has historically focused on hosts with highly stable, specific, and functionally important microbiomes (1). For example, the obligate nutritional endosymbionts of sap- and blood-feeding insects, which exemplify this scenario, have long served as important model systems. More recently, however, wider microbial explorations have shown that this kind of association does not apply to all insect taxa. In some insect groups, microbiomes are more variable, and the constituent microbes have low or no specificity to their hosts (2, 3). Moreover, the functional importance of these microbiomes is often unclear—and possibly variable, low, or nonexistent in some groups (4). Understanding the nature of microbiomes in these insects is not only important to their biology, but also for a broader understanding of insect microbiome evolution.

The Lepidoptera (butterflies and moths) are a group in which microbiome form and function have been particularly difficult to resolve. Recent work indicates that larvae (caterpillars) typically harbor microbiomes that are variable, low-abundance, and transient (5–9). However, a crucial consideration for Lepidoptera and other insects that undergo complete metamorphosis is that microbial associations may change between life stages (10). In Lepidoptera, larvae and adults do often differ in the composition and absolute abundance of their microbiomes (6, 11–15). Beyond this pattern, however, the ecology of adult-stage microbiomes remains poorly understood.

There are three major unknowns about microbiomes of adult Lepidoptera that we sought to address in this study. First, it is unclear how variable microbiomes are among host individuals and taxa, and which ecological or evolutionary factors (e.g., host phylogeny) are associated with microbiome variation. Understanding these patterns could give insights into whether microbiomes are important for certain ecological traits, such as feeding, and into the likelihood of host-microbe coevolution. Second, it is unclear whether butterfly-associated microbes have any specificity to their host species or to higher taxonomic levels; tight specificity is a hallmark of obligate insect-microbe symbioses (3). Previous studies of adult Lepidoptera microbiomes have largely used 97% operational taxonomic units (OTUs) based on short fragments of the 16S rRNA gene. OTUs can encompass substantial strain-level diversity (16), hindering analyses of specificity. Third, the full diversity of eukaryotic microbes that may be associated with adult Lepidoptera has not been explored, as previous studies have exclusively focused on bacteria or fungi.

Our study focuses on the neotropical passion-vine butterflies (Nymphalidae: Heliconiini). This tribe, which includes the genus *Heliconius*, is a foundational system in evolutionary biology with over 150 years of scientific study (17). Given the scientific relevance of heliconiines (18), and the myriad roles that microbes can play in animal biology (19), it is important to know the structure and function of heliconiine-associated microbiomes. Heliconiine butterflies are also a potentially useful system for the field of host-microbe interactions, given the wealth of genomic data available (20) and the ability to rear and genetically manipulate members of this group (21). A handful of amplicon sequencing-based microbiome studies have included adult heliconiines, but these were limited to a small number of individuals and species (11, 22, 23). Consequently, we lack even basic information about the microbes harbored by heliconiine butterflies and how they may vary across individuals and the host phylogeny.

In addition to the practical advantages mentioned above, heliconiines are also an interesting system to address how host ecological traits evolve in concert with microbiomes. Within Heliconiini, the genus *Heliconius* exhibits unique feeding and life history traits relative to other genera: larvae feed on young passion-vine shoots and leaves as opposed to old foliage, adults of most species (besides *H. aoede*) feed on pollen as well as nectar, and they have distinctive defensive chemistry and a greatly extended adult-stage lifespan of several months (18, 24). These traits could influence and be influenced by microbiomes in many ways. For example, we hypothesized that pollen-feeding might be associated with a distinctive microbiome, either because it is a source of microbes (25) or of nutrients that shift resident microbiome composition (26), or, because the trait itself depends on metabolic contributions from novel microbes (11). A useful starting point to uncovering these kinds of interactions is to examine whether *Heliconius* have microbiomes that are internally consistent yet distinct from those of other heliconiine genera. If so, then we can zero in on the specific ecological and evolutionary drivers of variation, and the potential functional roles of microbiomes in *Heliconius* biology.

Here we addressed the following main questions: *i*) How variable are heliconiine microbiomes among host individuals, species, genera, and across the heliconiine phylogeny? *ii*) Are host traits, such as pollen-feeding, associated with microbiome variation? *iii*) How host-specific are butterfly-associated microbes? At the same time, we sought to facilitate future microbiome research on heliconiine butterflies by answering some additional questions: *iv*) How are microbes distributed within the butterfly body? *v*) How abundant are these microbes? *vi*) How reliable is amplicon sequencing of 16S rRNA genes for studying butterfly-associated bacterial communities, and *vii*) are there eukaryotic microbes we might be missing with targeted amplicon sequencing?

We collected 214 wild adult butterflies, representing 23 species and subspecies of Heliconiini, and characterized their microbiomes with 16S rRNA gene sequencing. While many adult Lepidoptera microbiome studies have used reared, captive specimens (6, 12, 13, 22, 27), these may give a biased picture of microbial community structure in the wild (11). We also used shotgun metagenomic sequencing on a subset of butterflies both to provide an untargeted assessment of microbial diversity (including eukaryotes) and to provide a finer-resolution picture of microbial specificity than is available from amplicon sequencing of short 16S rRNA gene regions. Furthermore, we used quantitative PCR to estimate the absolute abundance of bacteria across different heliconiine taxa and tissue types. Our work illustrates ecological and evolutionary dynamics of microbiomes in adult heliconiine butterflies and advances our general understanding of Lepidoptera-microbe interactions.

## Materials and Methods

### Field collections

The wild adult butterflies used for whole-body microbiome sequencing were collected from seven locations in Panama and Ecuador in May-August 2014 (more detail is provided in the supplemental file “Collection_localities.txt”). Butterflies were euthanized with ethyl acetate and stored in DMSO after removal of wings, following (28). We also stored two DMSO-only blanks to use as negative controls. In June 2016, we collected additional adult butterflies for gut and head/thorax sequencing from Gamboa and Pipeline Road, Panama. For these specimens, we dissected the gut (hindgut, midgut, and the distal ~1/2 of the foregut) using sterilized tools prior to storage in DMSO. The whole head and thorax (including the proximal foregut) were stored separately. Species or subspecies were identified based on morphology. Butterflies were collected under permit # SC/A-7-11 from Panama’s Autoridad Nacional del Ambiente and # 005-13 IC-FAU-DNB/MA from Ecuador’s Ministerio del Ambiente.

### Sample processing, qPCR, PCR and sequencing

We removed whole bodies and head/thorax samples from DMSO and, after homogenization, used approximately 50 mg subsamples of homogenate for DNA extractions with the MoBio PowerSoil kit following the manufacturer’s instructions. We added entire guts directly to DNA extraction tubes, in which they were homogenized during the first bead-beating step of the protocol. Two DMSO blanks and 30 DNA extraction blanks were also processed in tandem with the butterfly samples and sequenced. For dissected gut and head/thorax samples, we estimated total bacterial abundance using quantitative PCR (qPCR) with 16S rRNA gene primers (515F/806R) following the protocol described in ref. (5).

PCR amplifications (515F/806R primers, V4 region) and 2 × 150 bp Illumina MiSeq sequencing of 16S rRNA genes followed standard Earth Microbiome Project protocols (dx.doi.org/10.17504/protocols.io.nuudeww). For gut and head/thorax samples of 29 butterfly individuals of nine species, we also attempted PCR amplification with primers that target the ITS gene region of fungi (29). However, amplification success with these fungal-specific primers (as assessed by gel electrophoresis) was very low, suggesting a lack of abundant fungal DNA that was later corroborated with the shotgun metagenomic data (see below). DNA extracts from a subset of 15 amplicon-sequenced gut samples were used for shotgun metagenomic sequencing following the approach described previously (30) with an input DNA concentration of 0.75 ng/ul, KAPA HiFi HotStart ReadyMix and bead cleanup with Ampure XP beads at a 0.9x ratio.

### Amplicon data processing

Amplicons from the 2014 whole-body samples and 2016 gut and head/thorax samples were sequenced on separate runs, demultiplexed using idemp (https://github.com/yhwu/idemp), and combined for further processing. Cutadapt (31) was used to remove primer sequences. We then used the DADA2 pipeline (32) to quality-filter (max EE value = 1) and trim (150 bp forward, 140 bp reverse) reads, infer exact sequence variants (ESVs), merge paired-end reads, and remove chimeras. We classified ESVs using the RDP Naive Bayesian Classifier algorithm (33) against the SILVA training set v. 132 (34).

Further data processing and analyses were conducted in R v. 3.6.0 (35). We used decontam (36) for prevalence-based identification of putative contaminant ESVs based on 34 negative controls (DMSO and DNA extraction blanks and PCR no-template controls). The median percentage of contaminant sequences across butterfly samples was 0.09%, but two samples had >10% contaminants and were removed from further analysis. ESVs with < 100 total sequences across all samples (out of a combined total of 5.6 million sequences) were removed, as were ESVs classified as mitochondria or chloroplast, or bacteria lacking sub-domain identification. These ESVs combined typically made up a low proportion of reads from the libraries (median of 2.6% across all samples). We also used these data to correct qPCR-based absolute abundances for non-bacterial amplification. Specifically, we multiplied the proportion of bacteria in sequence libraries by the total number of 16S rRNA gene copies to obtain estimates of bacterial absolute abundance.

As the 16S rRNA gene amplicon libraries were highly variable in read depth across samples, we rarefied all samples to 5,000 sequences, filtering out 13 samples with lower sequence depth. We relabeled *Pantoea*, *Erwinia*, *Kluyvera*, *Citrobacter*, *Klebsiella*, and *Cronobacter* taxonomic assignments to *Enterobacter*. This step was taken as genera within Enterobacteriaceae are often polyphyletic and are difficult to resolve from short 16S rRNA gene regions (e.g., (37)), and we wanted to avoid spurious separation of ESVs among genera and resulting idiosyncrasies in genus distributions across butterflies. Another case of ambiguous taxonomy occurred with sequences classified as *Pseudomonas* by the SILVA reference database, which we label below as “*Entomomonas/Pseudomonas*”. *Entomomonas* is a recently described genus of apparently insect-specialized Pseudomonadaceae and includes several sequences retrieved from *Heliconius* genome assemblies (38).

### Amplicon and qPCR data analysis

Beta diversity statistics and plots are based on Bray-Curtis dissimilarities and/or UniFrac distances (weighted and unweighted). To obtain a bacterial phylogeny for the latter, we used the fragment-insertion method (39) to place our ESV sequences into the Greengenes reference tree (40). The butterfly phylogeny is from (41) (TreeBASE #Tr77496). Four of the butterfly species in our sample set contained specimens from two distinct subspecies (e.g., *Heliconius sara sara* and *Heliconius sara magdalena*). To include these in the species-level host phylogeny, we inserted subspecies tips halfway along the terminal branches to their sister subspecies. Hereafter we refer to these subspecies as “species” for simplicity.

To test for host-phylogenetic signal in microbiomes, we used Mantel tests with 9999 permutations to calculate the correlation between microbial community dissimilarities/distances and host phylogenetic distances (42). Intraspecific variation in microbiomes was handled by averaging the pairwise dissimilarities/distances between all individuals of one species and all individuals of another species. We used the phytools package (43) to visualize concordance between topologies of the host phylogeny and a dendrogram of bacterial community dissimilarities. Nodes were rotated with the “cophylo” function in phytools to maximize tip matching between the two trees.

Differences in overall community composition between host genera were tested with PERMANOVA as implemented in the vegan package (44). Using the “betadisper” function we corroborated that significant test results were due to host genus-level differences in location and not dispersion (45). We used a nonparametric statistical test (Wilcoxon rank-sum) to identify bacterial genera that differed in relative abundance between host taxa or between sample types (gut versus head/thorax) and applied a false discovery rate correction to the resulting p values.

We tested whether total bacterial abundance differed between *Heliconius* and other butterfly genera using qPCR data from gut samples and head/thorax samples. Each sample type was analyzed separately using log-transformed counts of bacterial 16S rRNA gene copies. We verified that residuals were approximately normally distributed and used a linear mixed-effects model as implemented in the nlme package (46), treating host genus (*Heliconius* vs. others) as a fixed effect and host species as a random effect.

### Metagenome data processing and analysis

For 15 gut samples, we obtained shotgun metagenomic data to complement the bacterial 16S rRNA gene amplicon dataset. We quality-filtered these reads with sickle (47) and trimmed adapters with cutadapt (31). We then used Bowtie 2 (48) to filter out reads matching a given sample’s corresponding host species’ genome, obtained from Lepbase (20). The two *Dryadula phaetusa* metagenomes were mapped to a genome of the sister species *Dryas iulia* as no *Dryadula* genome was available. Since there was a high proportion of host-derived reads, we focused here on describing microbial diversity using ribosomal RNA gene reads present in the metagenomes. With the host-filtered reads, we used phyloFlash (49) to find and classify eukaryotic and bacterial SSU rRNA reads. Bacterial community composition was compared between the amplicon and shotgun metagenomic datasets using a Mantel test. We also used phyloFlash to assemble 16S rRNA genes from the 150 bp shotgun reads. These longer sequences allowed us to estimate the phylogeny of *Orbus*, the dominant bacterium in these 15 samples. *Orbus* sequences were aligned with MUSCLE (50), curated with Gblocks (51), and used for maximum likelihood reconstruction with the phylogeny.fr implementation (52) of PhyML (53).

### Data availability

Amplicon data, metadata, and R code are available from figshare (figshare.com/projects/Heliconius_butterfly_microbiomes/70520). Metagenomes are available from MG-RAST (project no. MGP89563).

## Results

Adult heliconiine butterflies host whole-body microbiomes that are typically low in diversity and evenness. Across all individuals, the median ESV richness was 26, of which only 11 accounted for 95% or more of the sequences. The composition of these communities is also reasonably consistent within host species. For example, within our most deeply sampled population (*H. erato demophoon* in Gamboa, Panama; N = 23), a median of 84% of the 16S rRNA gene reads obtained from a given individual’s microbiome belonged to a core set of 10 bacterial genera (Fig. S1). Some of these genera are present in roughly similar relative abundances across individuals and across our two sampling years (Fig. S1). Applying the aforementioned calculation to all heliconiine species from which five or more individuals were sequenced, we found that this intraspecific consistency extends beyond *H. erato demophoon*. For most host species, over 80% of the constituent individuals’ microbiome reads belonged to the 10 most abundant bacterial genera for that species (median = 88%, range = 77-94%).

Microbiomes from whole, homogenized butterfly bodies are mainly composed of gut-associated taxa. Isolated guts are similar to conspecific whole-body microbiomes in their bacterial community profiles (Fig. S1), with many bacterial genera occurring at similar relative abundances in guts and head/thorax tissue (Table S1). Some bacteria did differ between whole-body and gut samples, such as *Wolbachia*, which usually inhabit reproductive tissues (54) (Fig. S1). Likewise, in both *Heliconius* and other heliconiine genera, the relative abundance of *Acinetobacter* was ~10-fold higher in head/thorax samples as compared with guts (Table S1). In contrast, *Orbus*, *Enterobacter*, *Asaia*, and some other dominant bacterial genera are enriched in gut tissue (Table S1).

We used metagenomes (N = 15) to examine the accuracy of amplicon sequencing for describing bacterial community composition in our broader sample set. Interindividual microbiome variation was highly correlated between the amplicon-based dataset and the metagenomic dataset (Mantel *r* = 0.63, p < 0.001). Furthermore, for six of the most abundant bacterial genera, relative abundances in amplicon libraries were highly predictive of relative abundances in metagenomes (Fig. S2). These results support the use of amplicon sequencing for evaluating bacterial community composition in adult heliconiine butterflies.

Adult *Heliconius* butterflies generally harbor a high absolute abundance of gut bacteria, though titers sometimes varied substantially even among conspecific individuals collected from the same location (Fig. 1A). On average, *Heliconius* harbored significantly higher (by ~5.5-fold) gut bacterial titers than species belonging to other passion-vine butterfly genera (p = 0.039). Bacterial communities associated with head and thorax tissue did not significantly differ in titer between *Heliconius* and other genera (p = 0.38; Fig. S3).

**Figure 1.**
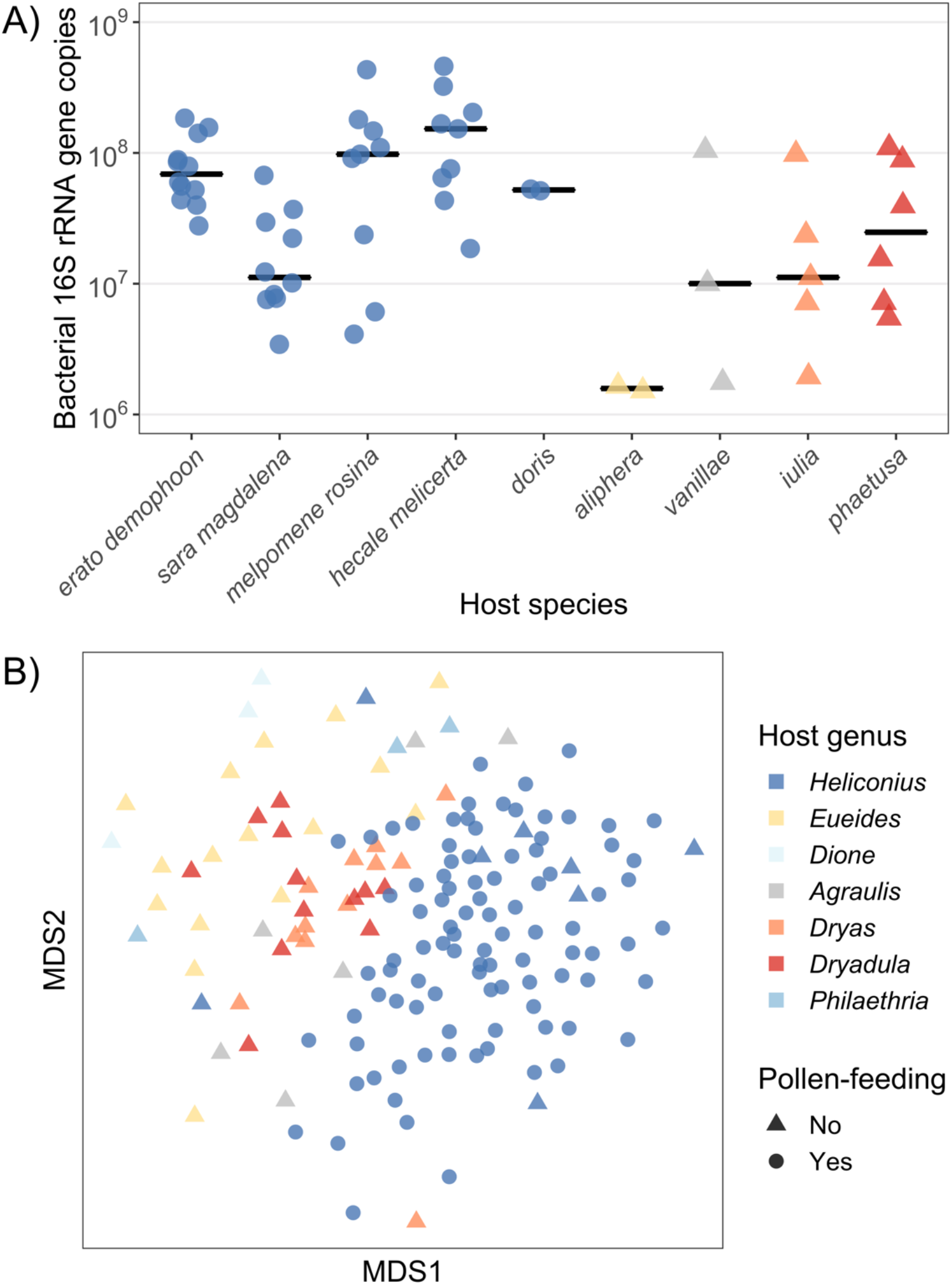
A) Adult *Heliconius* butterflies have high, though often variable, titers of gut bacteria. Shown are quantitative PCR-derived estimates of bacterial abundances in terms of the number of bacterial 16S rRNA gene copies per individual gut. These individuals were collected from Gamboa, Panama in 2016 (N = 42 *Heliconius*, 16 other Heliconiini). B) *Heliconius* host distinct adult-stage bacterial communities compared with related genera. Shown is an ordination of microbiome variation (Bray-Curtis dissimilarities) among all whole-body samples collected at various sites in Panama and Ecuador in 2014 (N = 104 *Heliconius*, 52 other Heliconiini). With the exception of *H. aoede* (dark blue triangles), pollen-feeding is exclusive to, and ubiquitous within *Heliconius*.

*Heliconius* also harbor compositionally distinct whole-body bacterial communities as compared with other passion-vine butterfly genera (Fig. 1B). Microbiomes clustered by host genera to varying degrees depending on the distance metric used. The effect was strongest with taxonomic or phylogenetic metrics that incorporate information on the relative abundances of exact sequence variants (ESVs) (Bray-Curtis: R^2^ = 0.13, p = 0.001; weighted UniFrac: R^2^ = 0.18, p = 0.001) as opposed to purely presence/absence-based metrics (Jaccard: R^2^ = 0.11, p = 0.001; unweighted UniFrac: R^2^ = 0.10, p = 0.001). This separation of *Heliconius* microbiomes from related butterfly genera is not fully explicable by adult-stage diet, as individuals of the non-pollen-feeding species *Heliconius aoede* mostly clustered with pollen-feeding *Heliconius* species (Fig. 1B). An analysis of species-by-species Bray-Curtis dissimilarities confirmed that *H. aoede* microbiomes are not uniquely distinct from those of pollen-feeding *Heliconius* species (Fig. S4).

As suggested by the host genus-level clustering, we found that variation in butterfly microbiome composition was associated with host relatedness (phylogenetic distance). More closely related butterfly lineages harbored more similar microbiomes (Bray-Curtis: Mantel *r* = 0.40, p = 0.01), and the topologies of the butterfly phylogeny and the dendrogram of microbiome similarity were moderately congruent (Fig. 2). This pattern was still statistically significant when microbiome variation was measured using weighted UniFrac distances (Mantel *r* = 0.31, p = 0.03), but not when using unweighted UniFrac (Mantel *r* = 0.03, p = 0.41) or Jaccard distances (Mantel *r* = 0.19, p = 0.08), which do not incorporate relative abundance information.

**Figure 2.**
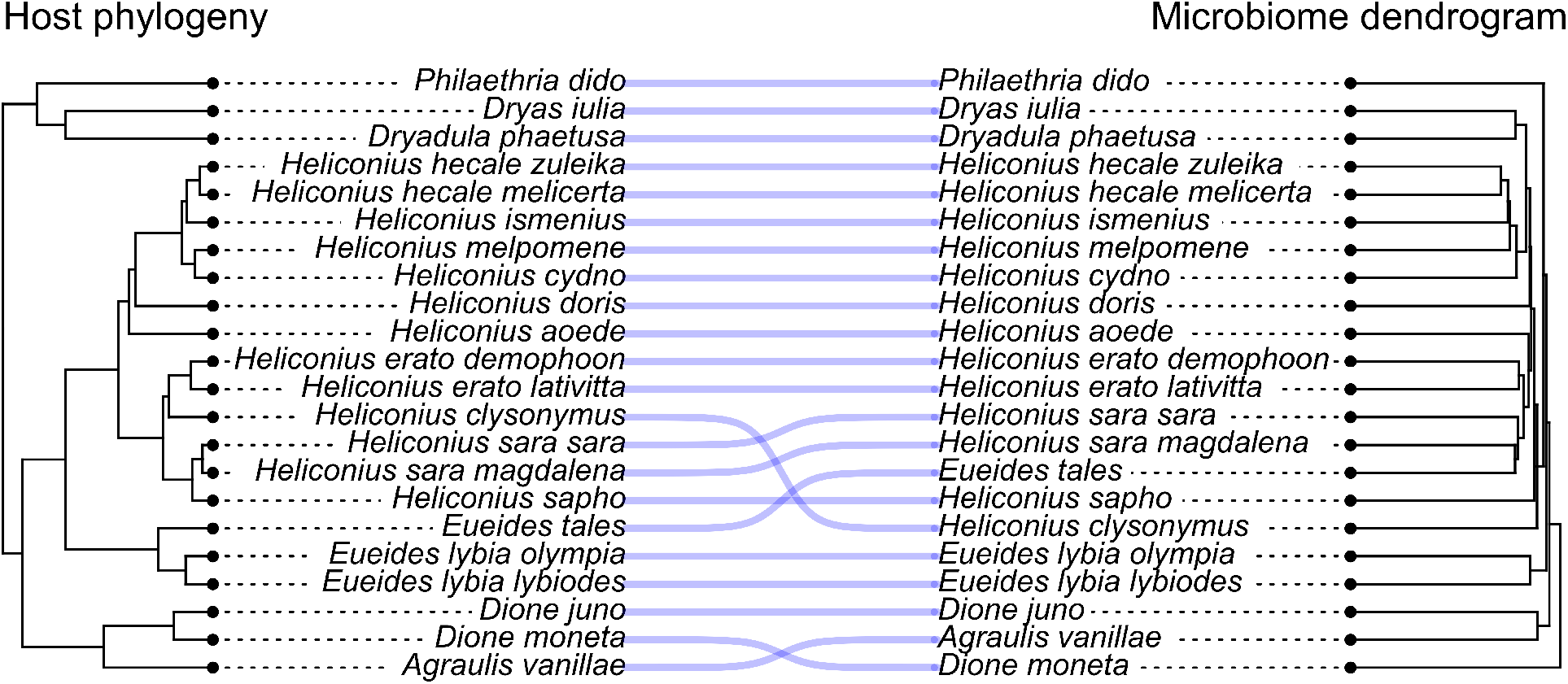
Variation in heliconiine butterfly microbiomes is correlated with host relatedness. Shown is the concordance between the host phylogeny (from (41)) and a dendrogram representation of microbiome variation among species (Bray-Curtis dissimilarities). Here, all nodes have been rotated to maximize tip matching. Note, however, that sets of parallel lines connecting tips between the phylogeny and the dendrogram do not always signify congruent branching structure.

We then determined which bacterial taxa contribute to the observed taxonomic (Fig. 1B) and phylogenetic (Fig. 2) structuring in overall community composition. The dominant bacterial genera *Enterobacter*, *Orbus*, and *Entomomonas*/*Pseudomonas* were proportionally more abundant in *Heliconius* versus the non-pollen-feeding butterfly genera, while *Asaia* and *Apibacter* showed the opposite pattern (Fig. 3). None of these bacterial genera, however, were exclusively restricted to one host feeding guild, genus, or species, and their relative abundances were occasionally highly variable (Fig. 3). We also analyzed the data at the ESV-level, including all individual ESVs that were reasonably abundant within at least one host species (≥ 5% within-species average). We did not find evidence for prevalent host species- or genus-restricted ESVs (Fig. 4). Most ESVs are present, albeit with varying relative abundances, across host genus and species boundaries.

**Figure 3.**
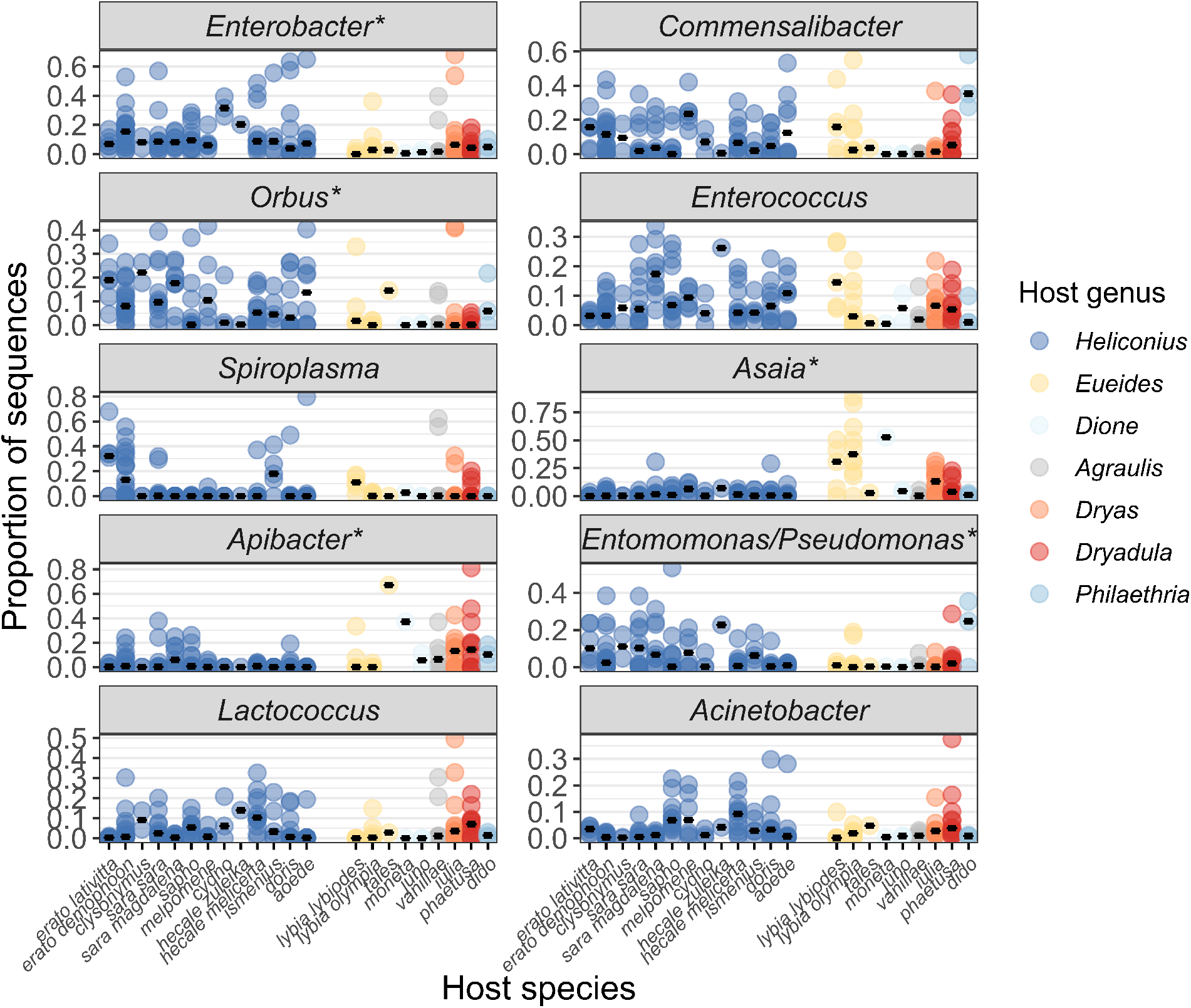
Dominant bacterial genera are largely shared among *Heliconius* and other butterflies, although some are differentially abundant. Shown are the relative abundances of the top 10 bacterial genera, ranked by mean abundance, in whole-body microbiomes. Dots indicate replicate individuals, and black bars indicate median proportions within a host species. Starred bacterial genera differed significantly in relative abundance between *Heliconius* and non-*Heliconius* butterflies (p < 0.05 after FDR correction). The arrangement of host species on the x axis corresponds to the phylogeny shown in Fig. 4. Note that *Enterobacter* here includes sequences originally assigned as *Klebsiella* and some other closely related Enterobacteriaceae genera (see Methods).

**Figure 4.**
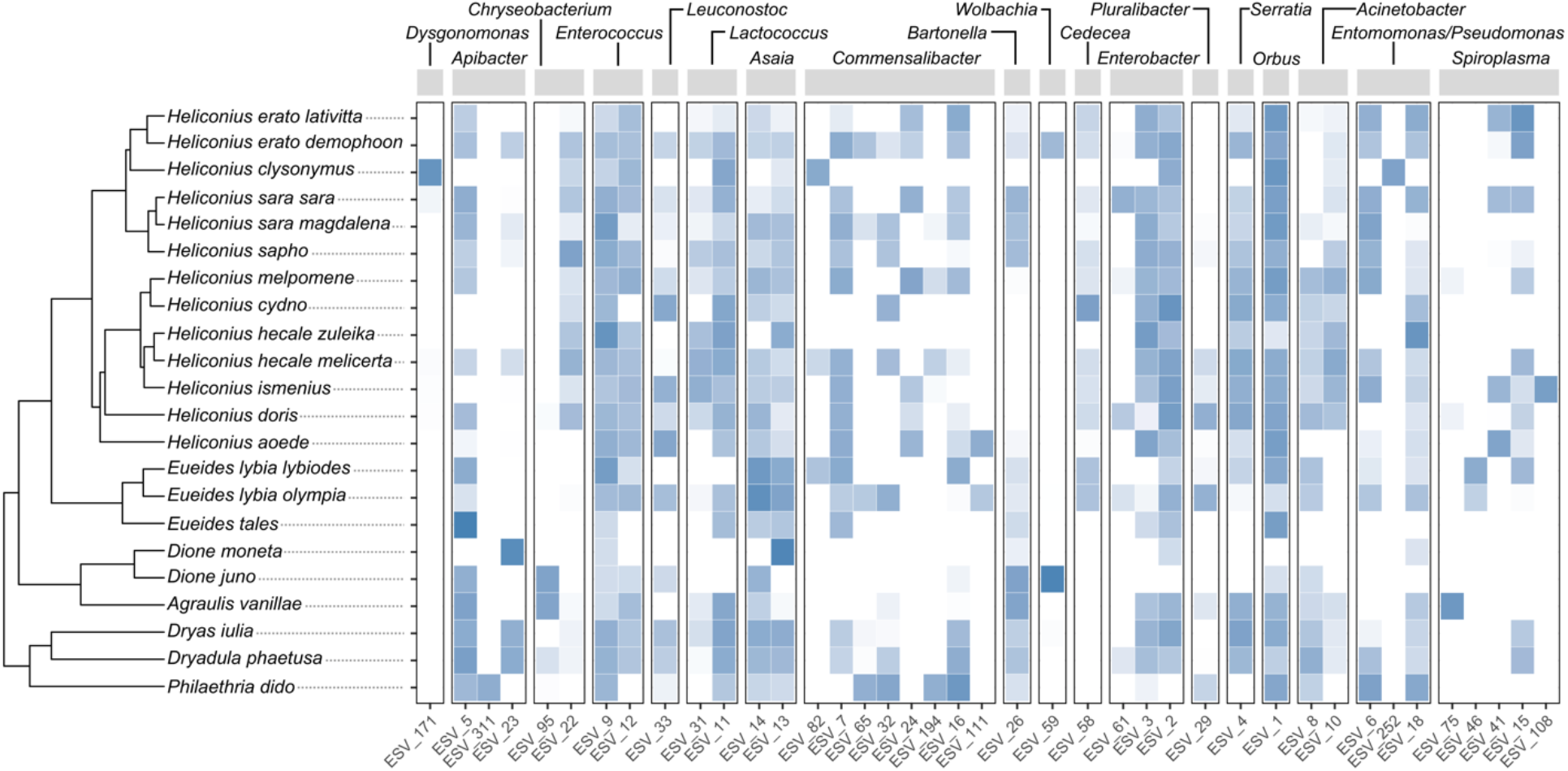
Most bacterial exact sequence variants (ESVs) are not host species- or host genus-specific. Shown are all ESVs in the dataset that had ≥ 5% mean relative abundance across conspecific individuals for one or more host species. For each ESV, the species-level mean relative abundance in whole-body samples, after log transformation, is indicated by the color of the cells (white = not detected in that species). Note that bacterial genera (labeled at the top) contained varying numbers of ESVs that met the aforementioned prevalence threshold.

Amplicon-derived ESVs are limited in their ability to resolve bacterial strains as they represent only a short region of the 16S rRNA gene (here, ~250 bp). To test for potential host specificity at a finer level of resolution, we obtained near-full-length 16S rRNA gene sequences from the bacterium *Orbus*, which is highly prevalent across gut and whole-body samples (Fig. 3, Fig. S1) and almost exclusively composed of a single ESV (Fig. 4). These sequences were reconstructed from the 12 metagenomes in which *Orbus* was sufficiently abundant. A phylogeny based on these sequences and other Orbaceae shows that butterfly-associated *Orbus* form a unique clade (Fig. 5). Host-phylogenetic or geographic structure was not evident within this clade. In fact, many of the Panamanian heliconiine butterflies harbored *Orbus* that have identical or near-identical 16S rRNA gene sequences to an *Orbus* strain isolated from an East Asian butterfly, *Sasakia charonda* (55).

**Figure 5.**
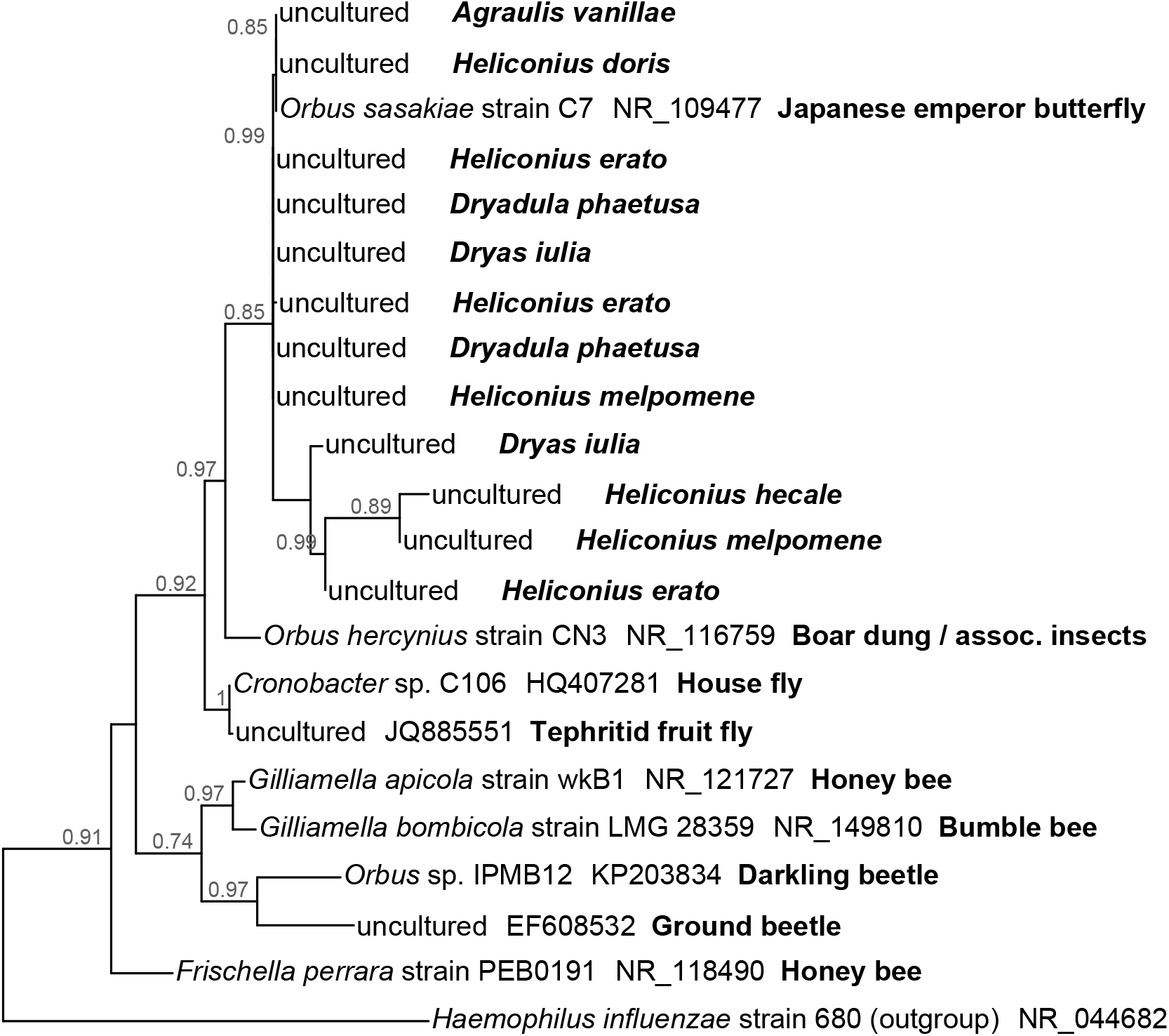
Maximum likelihood phylogenetic reconstruction of the bacterial family Orbaceae (Gammaproteobacteria: Pasteurellales). Some host taxonomic structure is apparent at the order level (i.e. Lepidoptera, beetles, bees, flies) but not within butterflies. 16S rRNA gene sequences from *Agraulis*, *Dryadula*, *Dryas*, and *Heliconius* were assembled from short metagenomic reads. Other 16S rRNA gene sequences are from GenBank. Branch support values are shown next to nodes. *Haemophilus influenzae* (Pasteurellaceae) was used as the outgroup.

We also used the shotgun metagenomes to search for microbial eukaryotes in butterfly gut samples and found that microsporidia and trypanosomatids were prevalent (27% and 47% of the 15 individual samples analyzed, respectively) (Fig. S5). Fungi and other microeukaryotic taxa were only sporadically detected, and at very low relative abundances. When trypanosomatids were detected in a given sample, we also sometimes detected 16S rRNA gene reads in the metagenomic data classified as *Kinetoplastibacterium*, an obligate bacterial endosymbiont of certain trypanosomatids (56). *Kinetoplastibacterium* was not detected without its corresponding trypanosomatid host (Fig. S5). This observation led us to reexamine the larger, amplicon-based bacterial dataset. We found that 9% of *Heliconius* individuals and 4% of individuals belonging to other heliconiine genera were infected with a trypanosomatid, as inferred by the presence of *Kinetoplastibacterium*. These proportions were not significantly different (Fisher’s Exact Test, p = 0.28). They are also likely underestimates, given the aforementioned prevalence of trypanosomatids among metagenomes (47%) and the potential for false negatives—in five metagenomes, trypanosomatids were detected without *Kinetoplastibacterium* (Fig. S5).

## Discussion

We found that adult-stage microbiomes in heliconiine butterflies are generally similar in composition among conspecific individuals and over time, and are also abundant within gut tissue (especially in *Heliconius*). Our estimates of total bacterial abundances (via qPCR), in parallel with those of ref. (23), support the hypothesis that bacteria actively colonize adult butterflies as opposed to passing through the gut transiently. These features contrast with the typical situation in lepidopteran larvae (5), reinforcing the idea that insect-microbe associations can differ strongly between life stages (10).

Although stable relative to larvae, dominant butterfly-associated microbes do exhibit higher levels of interindividual variability as compared with obligate nutritional endosymbionts or gut microbes of some other insect groups. For example, in honey bees, >98% of sequences belong to a honey bee-specific set of 5-9 bacterial species (57). We found that ~80-90% of sequences in a given butterfly species’ microbiome belonged to a set of 10 core bacterial genera. This contrast suggests that microbiomes are more facultative for butterflies or that there is greater functional redundancy among butterfly-associated microbes.

We also discovered that *Heliconius* butterfly microbiomes are distinct from those of other heliconiine genera, in terms of overall community structure (Fig. 1B), relative abundances of specific bacterial genera (Fig. 3), and total numbers of gut bacteria (Fig. 1A). This finding opens the possibility that one or more of the ecological traits specific to *Heliconius* influences, and may be influenced by, the microbiome. As these traits evolved in tandem (18) it is difficult to disentangle their potential links to microbes, but our analysis of the species *Heliconius aoede* suggests that pollen-feeding is not a primary driver. *H. aoede* does not pollen-feed (58, 59), yet its microbiomes were not uniquely distinct from pollen-feeding *Heliconius* species (Fig. 1B, Fig. S4). This result contrasts with previous work on butterflies finding a strong association between adult-stage feeding ecology (nectivory versus frugivory) and adult gut microbiomes (23).

Larval feeding ecology, however, warrants further investigation as a potential driver of variation in adult gut microbiomes. Microbiomes of butterfly larvae and adults are not fully decoupled: in *H. erato*, some adult-stage microbes appear to be carried over from the larval stage (11). All *Heliconius* (including *H. aoede*) feed on young passion-vine foliage while other heliconiine genera feed on mature foliage (18). Related variation in diet-derived microbes, or in host processes that determine which microbes persist through metamorphosis (10, 60), could potentially underlie variation in adult-stage microbiomes.

In addition to the clear separation of *Heliconius* from other heliconiine genera (Fig. 1B), there was also a strong signal of the host phylogeny in heliconiine butterfly microbiomes (Fig. 2). The strength of this signal was comparable to that observed in gut microbiomes of mammals (61) and other host groups (42). This result is also in agreement with a recent survey of neotropical butterflies, in which microbiomes were found to be phylogenetically structured across a broad diversity of hosts (six different families) (23). Thus, even in a comparatively young radiation such as the one studied here (41), host phylogenetic history is clearly an important factor shaping the composition of adult butterfly microbiomes. The question then becomes, how has this pattern (also known as phylosymbiosis (62)) arisen, and what does it signify?

There are at least two, non-mutually exclusive processes that can lead to phylosymbiosis: host-microbe co-diversification, and contemporary host filtering of environmental microbes (42, 63). In the former process, there is a shared evolutionary history of host and microbial lineages. This does not necessarily apply to the latter process, which occurs when host traits that influence microbial colonization, such as diet preference, are conserved to some degree across the host phylogeny. We suggest that co-diversification is unlikely in this system given the broad distribution of bacterial phylotypes across host species and genera (discussed below). Rather, host filtering of environmental microbes likely explains phylogenetic structure in heliconiine microbiomes, as is the case in a wide variety of other animal groups (42). A priority for future work on butterflies is to identify the specific traits underlying host filtering. In Heliconiini, two candidates are larval host plant use (i.e., *Passiflora* species identity and age of tissue consumed) and adult foraging behavior (especially whether pollen is collected, and from which plant species), both of which exhibit phylogenetic signal (18, 64, 65).

Bacterial community-level variation among butterfly genera and across the phylogeny was largely driven by shifts in the relative abundance of shared bacterial genera and ESVs, as opposed to the presence or absence of host-specific bacterial taxa (Figs. 3, 4). Moreover, a finer-resolution analysis of the nearly ubiquitous bacterium *Orbus* did not find evidence for specificity within butterflies (Fig. 5). This pattern is notable in part because it weighs against the co-diversification model described above, and in part because it provides a contrast to the high degree of host specificity documented in a number of other insect-microbe symbioses (66–69). While not highly host-specific, at least some of the dominant bacteria in butterflies appear to be insect specialists as opposed to cosmopolitan environmental taxa. For example, phylogenetic evidence supports a general insect association for *Entomomonas* (38), *Apibacter* (70), and *Orbus* (Fig. 5) (71). The ecology of these groups is poorly understood; one important unknown is how they are transmitted among butterflies, which may be via flowers (72) or other shared resources.

Shotgun metagenomes allowed us to corroborate the amplicon-based bacterial data (Fig. S2) and led to the discovery of putative eukaryotic parasites (trypanosomatids and microsporidia) in gut tissue of many adult heliconiine butterflies (Fig. S5). Beyond fungi (12, 23), which do not appear to be abundant in heliconiines, microeukaryotes are almost unknown from adult-stage Lepidoptera—with the exception of the neogregarine *Ophryocystis elektroscirrha* in monarch butterflies (73). Given their potential interactions with gut bacteria and relevance to host fitness, more targeted analyses of butterfly-associated microeukaryote diversity are clearly warranted.

While effects of endosymbionts such as *Wolbachia* and *Spiroplasma* have been documented (74), a major open question is what functional role gut microbes play in the biology of adult heliconiines and Lepidoptera generally (75). In *Heliconius*, our hypothesis that microbes mediate pollen-feeding is not strongly supported. To directly influence pollen digestion—which occurs extraorally using saliva exuded from the proboscis (76)—microbes would likely need to inhabit the proboscis or salivary gland. Yet we saw no clear signal of elevated microbial abundances or unique microbial taxa in these tissues in *Heliconius* as compared with non-pollen-feeding butterflies (Fig. S3). Endogenous pollen-digestion mechanisms (59, 77, 78) may be sufficient.

Other microbial roles in host nutrition are possible, although a recent experiment on the butterfly *Speyeria mormonia* did not find evidence for nutrition-related functions (79). We suggest that colonization resistance (i.e., protection from pathogens and parasites), which is a feature common to many symbioses (63, 80), could be a primary ecological function of adult butterfly gut microbiomes. We found that heliconiines are frequently infected by trypanosomatids and microsporidia, as well as *Serratia* (Fig. 4), which are common opportunistic bacterial pathogens in insects (81, 82). Adult butterflies may be similar to pyrrhocorid bugs, tsetse flies and social bees, in which gut bacteria provide an important layer of defense against related parasites and pathogens (83–87). Importantly, colonization resistance can readily evolve even in symbioses lacking strong host-microbe specificity (88), such as that between heliconiines and their adult-stage microbiomes.

## Conclusions

This study provides an in-depth characterization of *Heliconius* and other passion-vine butterfly microbiomes, adding a new dimension to a classic model system in evolutionary biology. The characteristics of adult-stage microbiomes we report contrast with larval Lepidoptera, emphasizing that holometabolous insects are able to interact with microbes in very different ways across life stages (10). However, many Lepidoptera do not feed or even have a digestive tract as adults (89), and we do not yet know whether patterns from heliconiines or other butterflies are generalizable to these other groups. We further suggest that heliconiines may serve as a useful system for exploring how animal ecology and life history relate to microbes.

*Heliconius* host phylogenetically structured microbiomes that are markedly distinct from those of their close relatives, and this finding sets the stage for experiments to test the specific host traits that may be involved. Finally, we uncovered novel microeukaryote diversity in butterflies, and hypothesize that they are parasites that interact antagonistically with gut bacteria. Further research on adult-stage microbiomes will help advance our understanding of both insect-microbiome evolution and of the biology of Lepidoptera, a diverse group of considerable ecological, societal, and scientific importance (89, 90).

## Acknowledgments

This material is based on work supported by the Cooperative Institute for Research in Environmental Sciences, the Ecology and Evolutionary Biology Department at the University of Colorado Boulder, a National Science Foundation Graduate Research Fellowship (1144083), and a postdoctoral fellowship from the USDA National Institute of Food and Agriculture (2018–08156), to TJH. This work was further supported by grants from the Simons Foundation (429440) and by funding from the Smithsonian Tropical Research Institute. We thank Jessica Henley for assistance with sequencing, the Smithsonian Tropical Research Institute staff for administrative support and the Panamanian and Ecuadorian Authorities for permission to collect butterflies. We also acknowledge the reviewers for helpful comments that improved the manuscript.

## Supplementary Figures

**Figure S1.**
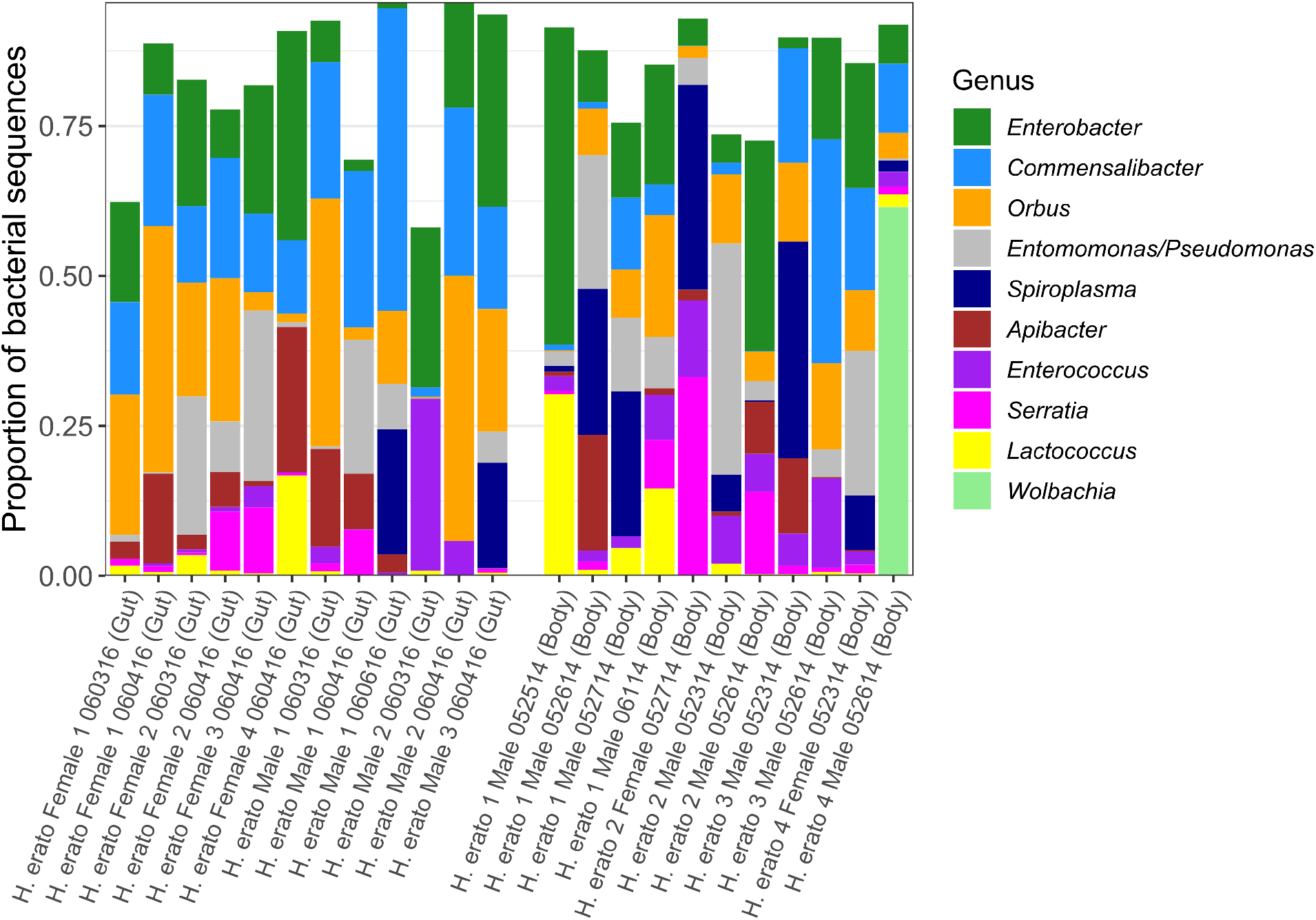
Whole-body bacterial communities are consistent among individuals and predominantly represent gut-associated taxa. Shown are the relative abundances of the top 10 bacterial genera (ranked by mean abundance) across *H. erato demophoon* collected from Gamboa, Panama. Remaining white space represents sequences belonging to other genera; these made up a median 16% of sequence libraries across individuals. Samples on the left are isolated guts from individuals collected in 2016, while samples on the right are whole-body homogenates from individuals collected in 2014. Note that *Enterobacter* here includes sequences originally assigned as *Klebsiella* and some other closely related Enterobacteriaceae genera (see Methods).

**Figure S2.**
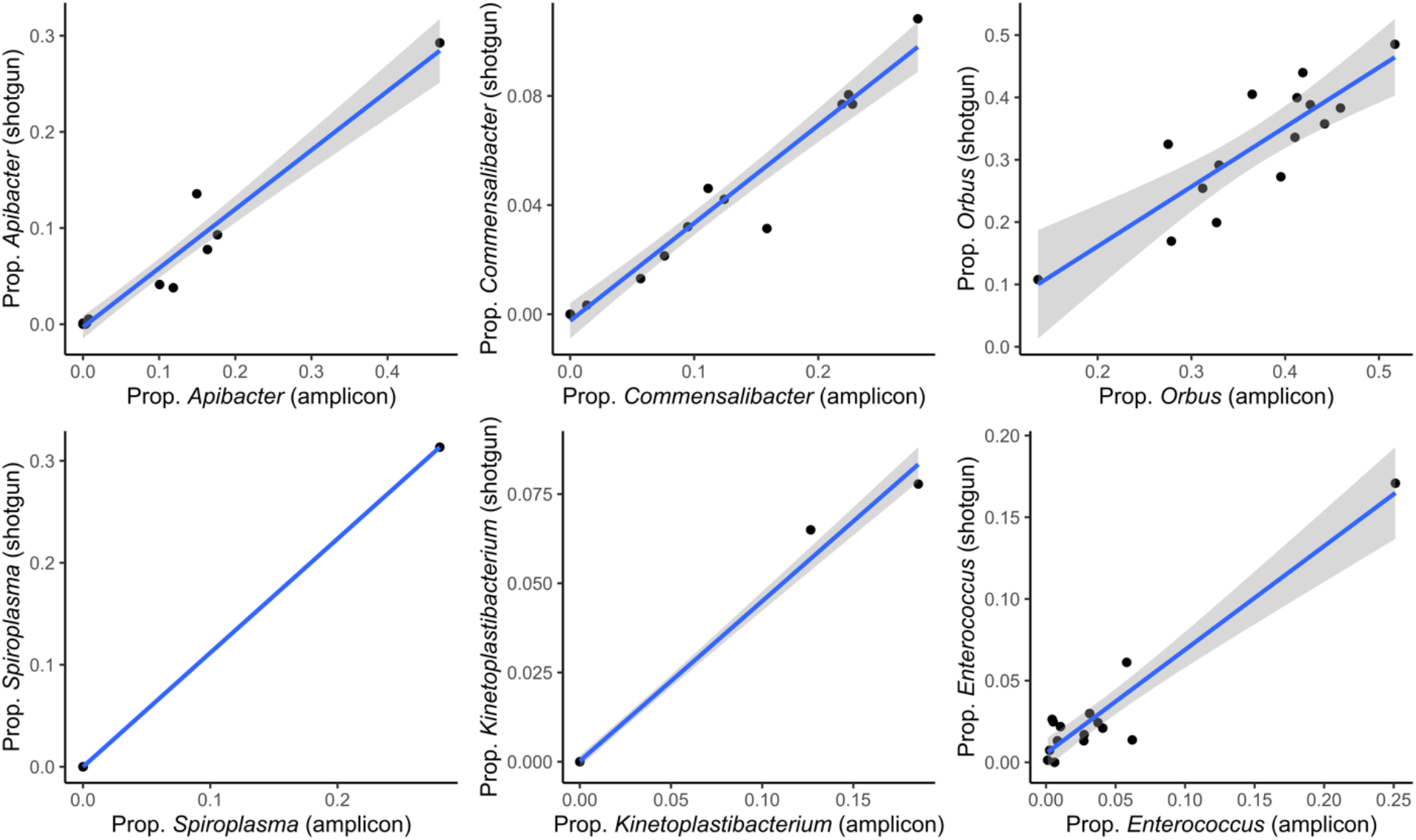
Bacterial genus-level relative abundances are highly correlated between the amplicon sequence libraries and metagenomes. Shown are relative abundances of six of the most abundant genera across the 15 butterfly gut samples for which we obtained both amplicon and shotgun metagenomic sequence data.

**Figure S3.**
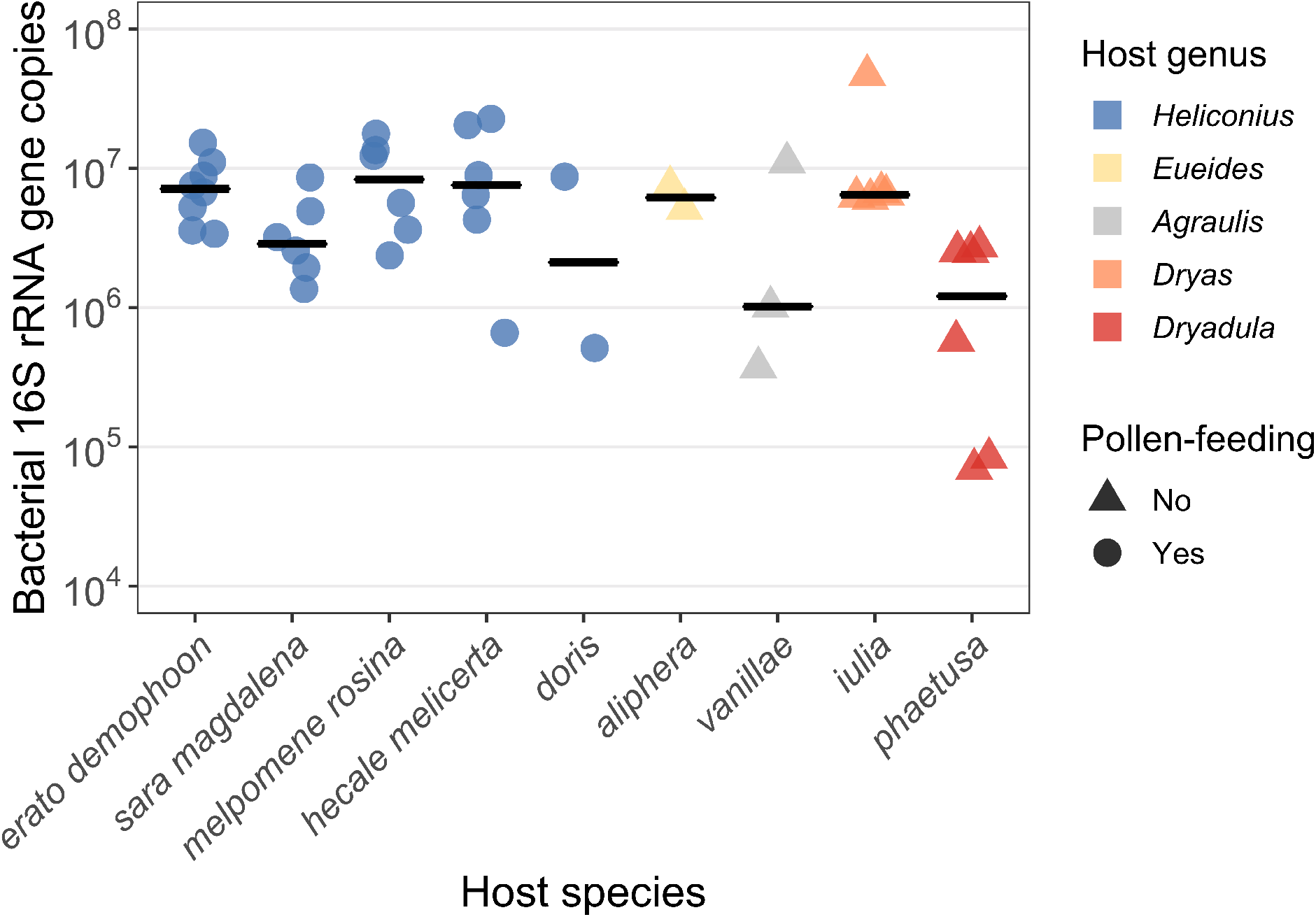
Absolute abundances of bacteria in the head and thorax of adult heliconiine butterflies, derived from qPCR. Shown are the number of bacterial 16S rRNA gene copies per subsample (approx. 50 mg) of homogenized, combined head and thorax tissue. These individuals were collected from Gamboa, Panama in 2016 (N = 28 *Heliconius*, 16 other Heliconiini).

**Figure S4.**
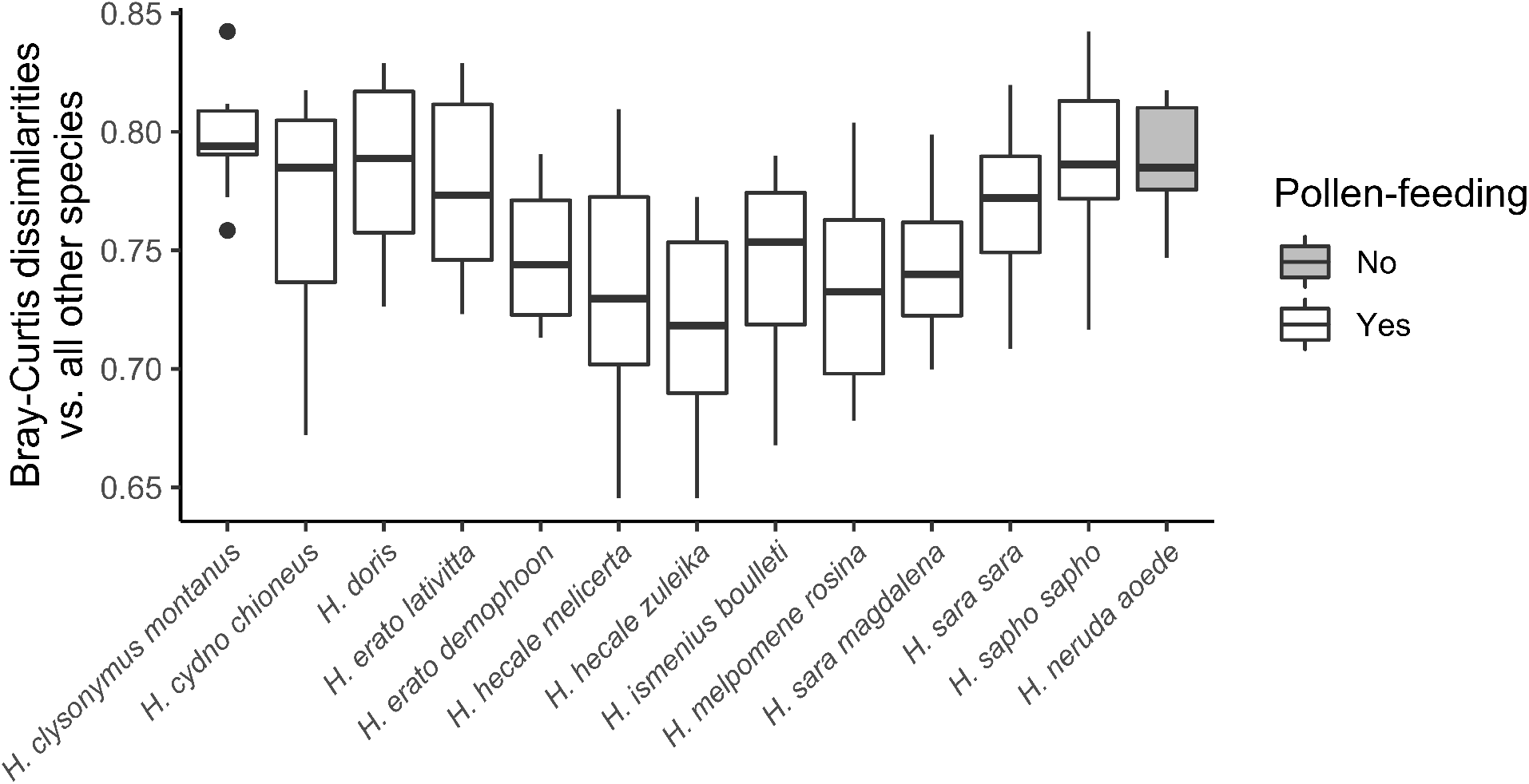
The non-pollen-feeding species *Heliconius aoede* is not uniquely distinct from pollen-feeding *Heliconius* species. Whole-body microbiomes from each focal species (x axis) are compared to all other species using Bray-Curtis dissimilarities. For a given comparison of two species’ microbiomes, intraspecific replication was handled by averaging dissimilarities among all pairs of individuals.

**Figure S5.**
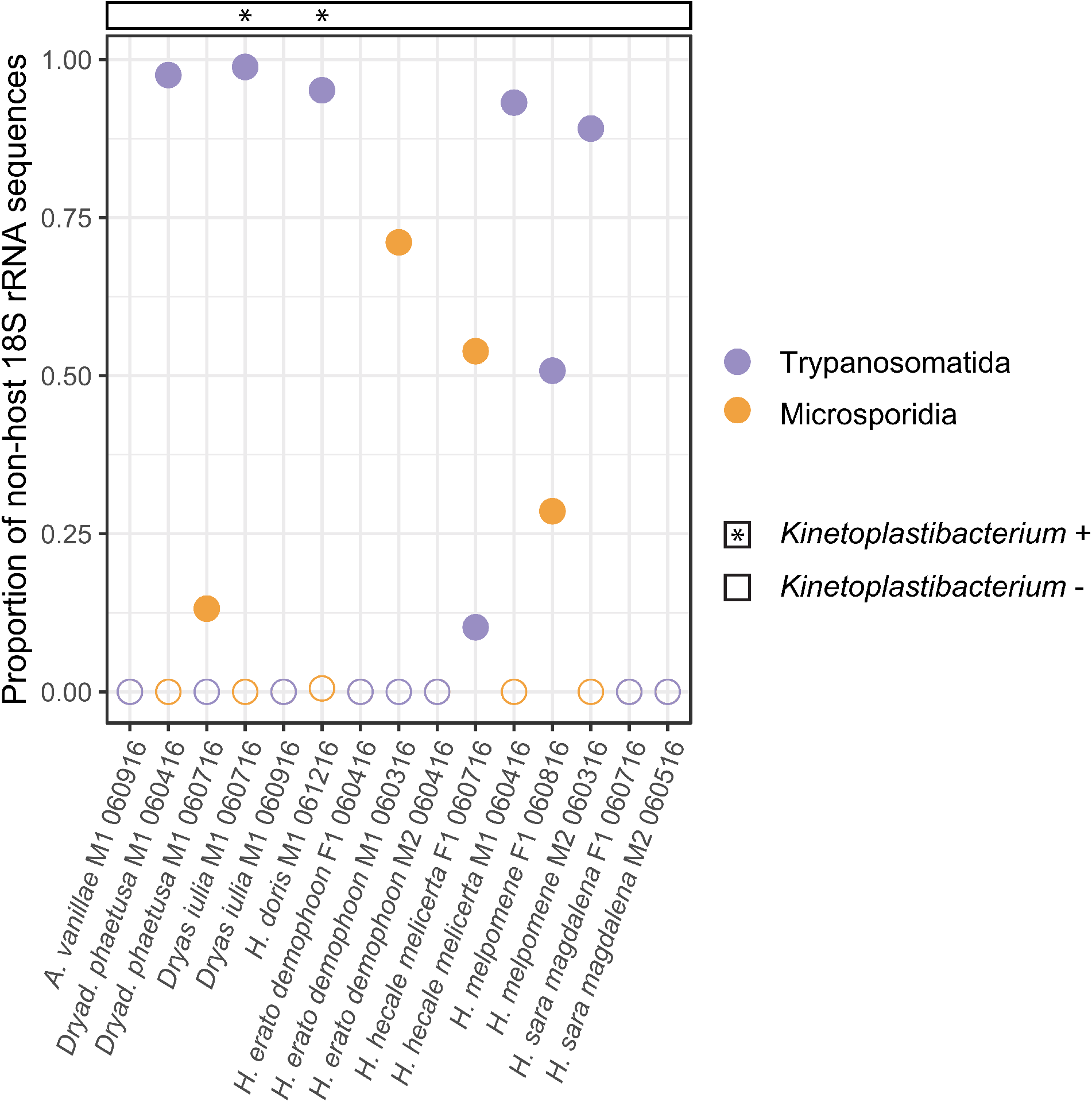
Microsporidia and Trypanosomatidae (related to *Crithidia*, *Leishmania*, and *Trypanosoma*) are prevalent in the 15 butterfly individuals with sequenced gut metagenomes. Points show the proportion of each microeukaryotic taxon out of all non-host 18S rRNA sequences identified by phyloFlash. Circles are filled if the proportion of microsporidia or trypanosomatids exceeded 0.01. Asterisks indicate samples in which *Kinetoplastibacterium*, a bacterial endosymbiont of trypanosomatids, was detected in metagenomes. Other microeukaryotes not shown here comprise a variety of very low-abundance taxa (i.e. ≤ 30 total reads per metagenome) classified mainly as coccidia, ascomycete fungi, acanthamoeba, and algae.

## Supplementary Table

**Table S1.**
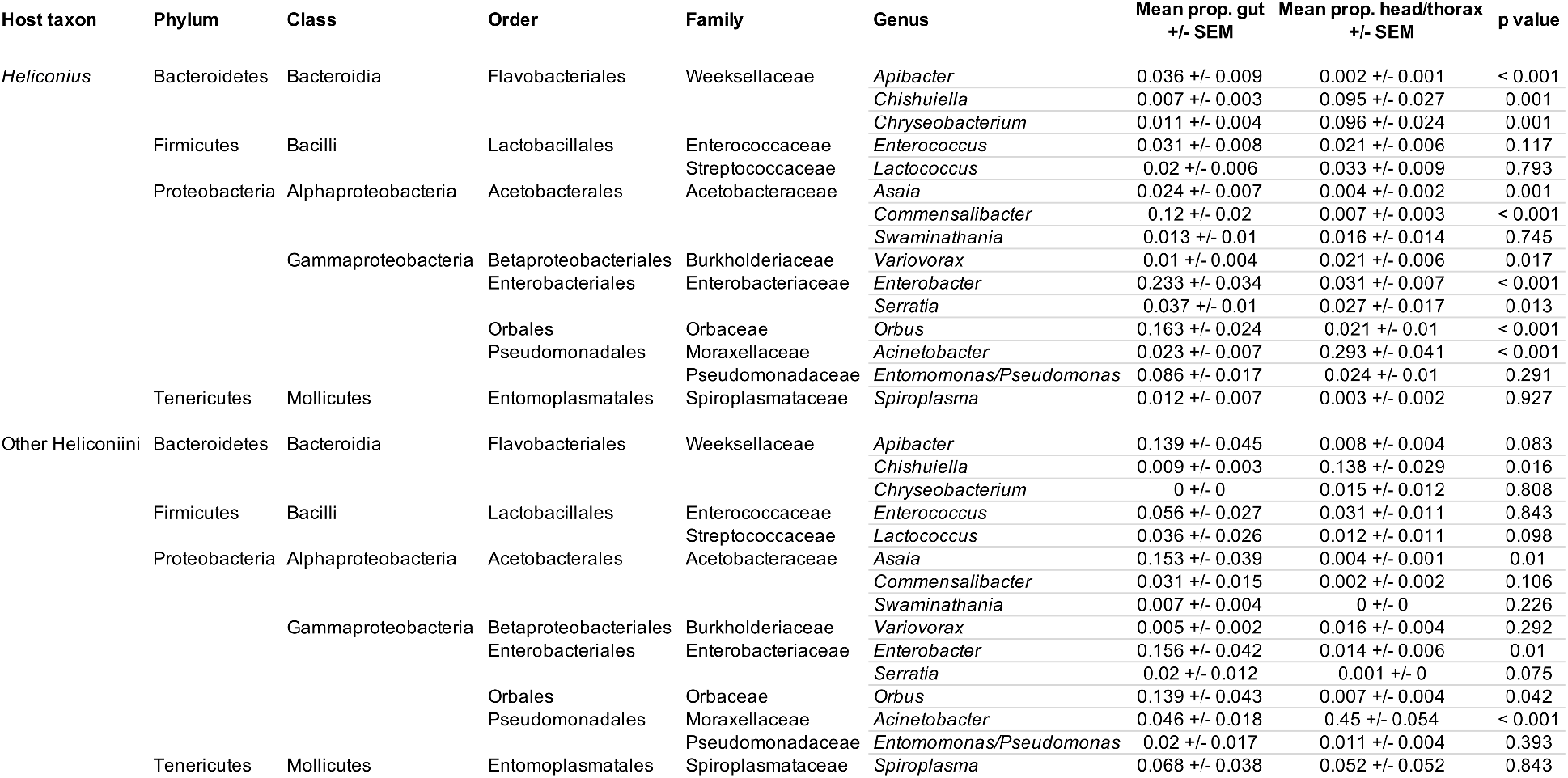
Relative abundances and within-body distribution of the top 15 bacterial genera (ranked by mean abundance) across butterflies collected from Gamboa, Panama in 2016. These individuals were dissected to compare microbiomes between isolated gut tissue and the combined head and thorax. Abundances are shown for *Heliconius* (top) and species belonging to other heliconiine genera (bottom) separately. P values are from a nonparametric statistical test of proportions in guts versus head/thorax samples, after FDR correction (see Methods).

